# Versatile Multicolor Nanodiamond Probes for Intracellular Imaging and Targeted Labeling

**DOI:** 10.1101/108720

**Authors:** Kerem Bray, Leonard Cheung, Igor Aharonovich, Stella M. Valenzuela, Olga Shimoni

**Affiliations:** School of Mathematical and Physical Sciences, Faculty of Science, University of Technology Sydney, Ultimo, NSW, 2007, Australia; School of Life Sciences, Faculty of Science, University of Technology Sydney, Ultimo, NSW, 2007, Australia; Institute of Biomedical Materials and Devices (IBMD), University of Technology Sydney, Ultimo, NSW, 2007, Australia

**Keywords:** nanodiamonds, bio-imaging, fluorescent probes, silicon-vacancy centers

## Abstract

Diamond nanoparticles that host bright luminescent centers are attracting attention for applications in bio-labeling and bio-sensing. Beyond their unsurpassed photostability, diamond can host multiple color centers, from the blue to the near infra-red spectral range. While nanodiamonds hosting nitrogen vacancy defects have been widely employed as bio-imaging probes, production and fabrication of nanodiamonds with other color centers is a challenge. In this work, a large scale production of fluorescent nanodiamonds (FNDs) containing a near infrared (NIR) color center – namely the silicon vacancy (SiV) defect, is reported. More importantly, a concept of application of different color centers for multi-color bio-imaging to investigate intercellular processes is demonstrated. Furthermore, two types of FNDs within cells can be easily resolved by their specific spectral properties, where data shows that SiV FNDs initially dispersed throughout the cell interior while NV FNDs localized in a close proximity to nucleus. The reported results are the first demonstration of multi-color labeling with FNDs that can pave the way for the wide-ranging use of FNDs in applications, including bio-sensing, bio-imaging and drug delivery applications.

---

Targeted sub-cellular imaging is crucially important for understanding biological processes and developing new drugs. To this extent, fluorescent probes are sought after for a range of potential applications, from mapping cellular environments, measuring cell temperatures and amino acids concentrations to monitoring drug localization within the body.^[1, 2]^ Despite numerous advantages, majority of fluorescent probes, such as quantum dots or fluorescent proteins, exhibits undesirable properties, including blinking, photo-bleaching and/or toxicity, limiting their use for biological imaging.^[3, 4]^

On the other hand, fluorescent nanodiamond (FNDs) particles are a promising material for bio-sensing and bio-imaging due to their ability to host a variety of bright, photostable color centers originated from atomic defects in the carbon lattice.^[5, 6]^ Furthermore, FNDs possess inherent biocompatibility and high surface tunability.^[7-9]^ To date, there is abundance of research on the application of FNDs containing nitrogen-vacancy (NV) defects for bio-imaging and drug delivery.^[10-17]^ However, NV FNDs have excitation and emission in visible range that can contribute to tissue absorption and auto-fluorescence. Therefore, FNDs containing centers with narrowband emission at the near IR (NIR) spectral range are particularly advantageous to achieve a better signal to noise ratio imaging for long term cellular imaging.^[18]^ In addition, excitation of NIR emitters can be achieved using a longer wavelength, therefore minimizing tissue light absorption. Finally, use of NIR luminescent probes is valuable in bio-imaging as it presents a greater tissue penetration depth comparing to visible range fluorophores.^[18]^

One particular defect that meets these criteria is the silicon vacancy (SiV) color center in diamond that has narrowband emission at 738 nm. Despite clear advantages of SiV ND properties, reliable fabrication and application of SiV FNDs for sustained biological applications remains a pressing issue and challenge.^[19]^

In our work, we report on a scalable approach to produce FNDs hosting the SiV emitters suitable for bio-imaging. We confirm that the fabricated nanoparticles hold NIR fluorescent properties, free from contamination and with sizes compatible for bio-imaging. We further demonstrate the concept of utilization of FNDs as multi-color staining for intercellular immunofluorescence. In fact, we show simultaneous and targeted labeling of different regions of cell interior using two different types of FNDs, such as containing SiV and NV defects. The targeting has been achieved due to the fact that the surface of nanodiamonds can be easily modified to attach targeting biomolecules using standard chemical procedures. Furthermore, we can effectively resolve two types of FNDs within cells by their specific spectral properties, where we find that SiV FNDs dispersed throughout the cell interior and NV FNDs localized in a close proximity to nucleus. Finally, we show long term imaging of FNDs up to 5 hours, where timed uptake of FNDs show cell induced mechanism of FNDs aggregation. Our results are the first demonstration of multi-color labeling with FNDs that can pave the way for the wide-ranging use of FNDs in applications, including bio-sensing, bio-imaging and drug delivery applications.

The SiV containing FNDs have been produced via bead-assisted sonication disintegration (BASD) process,^[20, 21]^ where chemical vapor deposition (CVD) grown diamond polycrystalline thin films were crushed into nanoparticles. The flow of the fabrication process is shown in **Figure 1**. To unambiguously show that the fabricated particles with SiV are made of diamond, Raman spectroscopy was employed to determine the characteristic diamond peak at 1332 cm^-1^ (**Figure S1**). To prove that the diamond particles host bright SiV color centers, the dispersion was characterized using a home built scanning confocal microscope using a 532 nm excitation source through a high numerical aperture (NA = 0.9) objective at room temperature. FNDs with NV centers were pre-characterized using the same system. **Figure 2a** shows the photoluminescence spectra, where we observed the characteristic zero phonon line (ZPL) at 737 nm and 640 nm for the SiV (red) and NV (blue) color centers, respectively. NV FNDs produce a broad red fluorescence ranging from 640 to 730 nm, while SiV FNDs show a sharp narrow peak ranging between 735 to 760 nm. It can be clearly seen that these two types of FNDs can be distinguished using their spectral characteristics.

**Figure 1.**
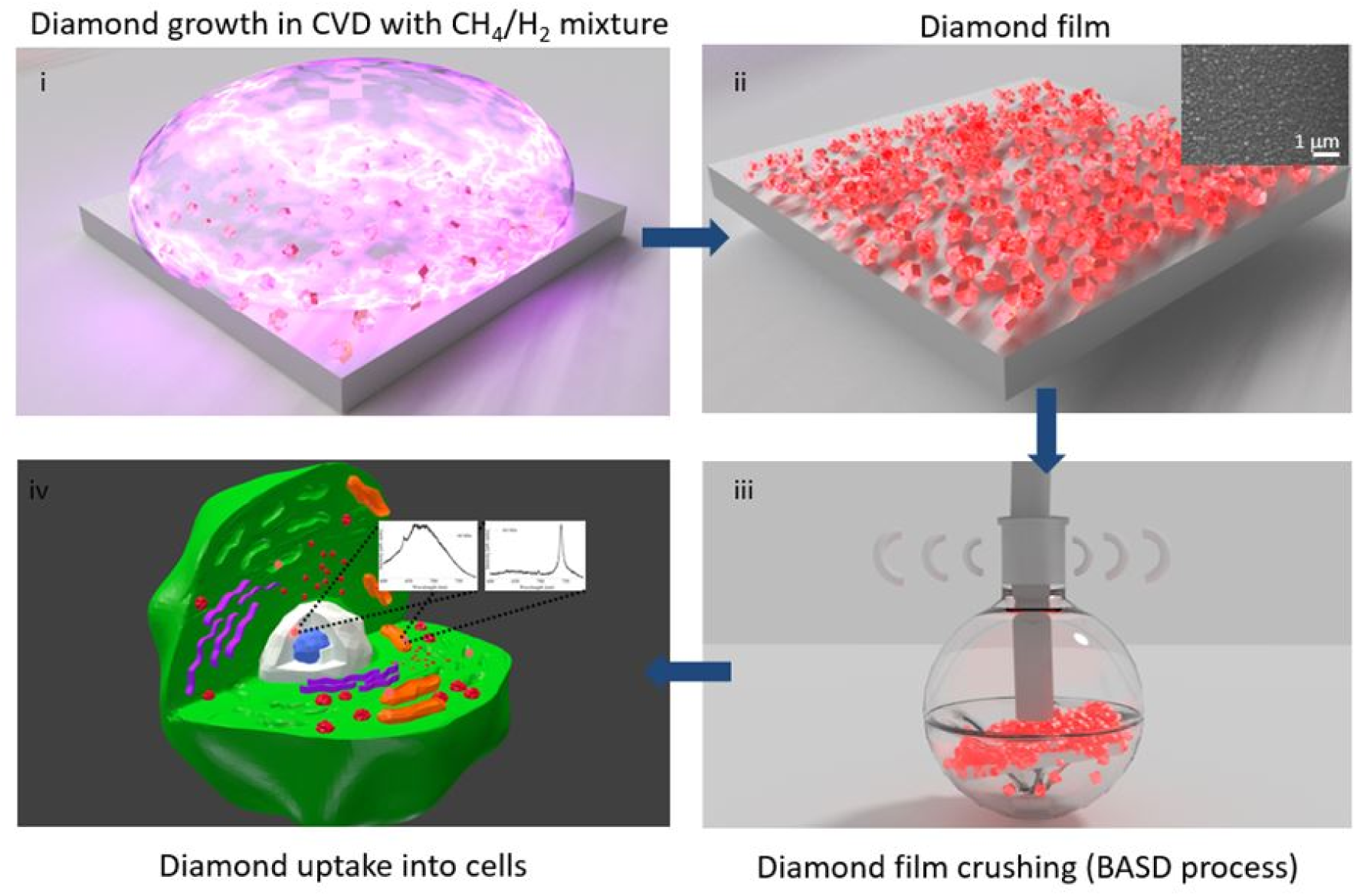
Synthesis of SiV containing FNDs (i). Silicon substrate seeded with detonation nanodiamond (4-5 nm) is subjected to CVD plasma using a standard hydrogen/methane gas mixture to produce highly dense polycrystalline diamond thin film (ii). SEM micrograph (inset) shows diamond thin film of SiV containing FNDs crystals. The CVD diamond film was isolated and was subjected to the BASD process (iii). To remove residual particles and graphite, the diamond solution then underwent strong acid reflux for 24 hours, washed through centrifugation with water to produce SiV FNDs in solution for intercellular bio-imaging (iv).

**Figure 2.**
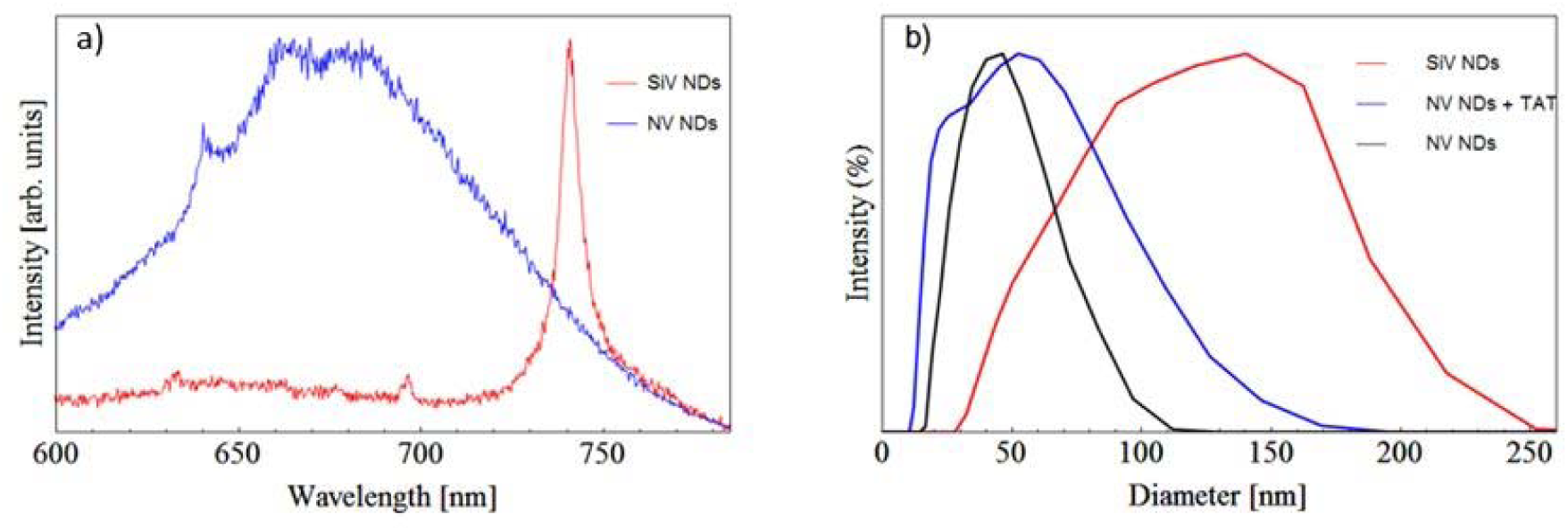
Characterization of different types of FNDs. (a) Typical room temperature spectra of SiV (red) and NV (blue) color centers under a 532 nm excitation source. (b) Particle size distribution by number of SiV (red), NV (black) and NV containing FNDs with TAT (blue) showing sizes <150 nm in diameter.

The size and zeta potential of FNDs were characterized by Zetasizer Nano ZS (Malvern Instruments Ltd) to test its suitability for bio-applications (Figure 2b). The measurements of size and zeta potential show the FNDs to be 141.1±49.4 nm and 44.8±13.7 nm in diameter with a zeta potential of -7.4±3.7 mV and -45.4±21.2 mV for the diamonds containing SiV and NV color centers, respectively (Table S1). The origin of negatively charged zeta potential is due to oxidation treatments that FNDs were subjected during the purification processes that produced oxygen-containing groups, such as carboxylic, hydroxyl or ester groups. The results confirm that the ND samples are relatively small, stable and dispersed in aqueous solution that is suitable for bio-applications.

Once we established the robust fabrication of the SiV containing FNDs, we precede their use in bio-labeling. A particular goal in our work is to demonstrate multiple, distinct cellular or molecular targets specifically through the conjugation of antibodies, drugs or organic chemicals. To this end, we utilize the convenience of carbon surface functionality to attach functional biomolecules to achieve biologically active nanoprobes. Specifically, FNDs containing NV defects were conjugated with trans-activating transcriptional activator (TAT, China Peptides Ltd) peptides using carbodiimide chemistry. TAT peptide is a small basic cell penetrating peptide (CPP) derived from HIV-1,^[22]^ and is known for successful cell membrane penetrating functionality that is effective at delivering materials of varying sizes from small particles to proteins into primary and other cells.^[23-25]^ The successful conjugation was monitored by surface charge change from -45.4±21.2 mV to +32.8±5.6 mV due to multiple positively charged lysine residues on the peptide chain. On the other hand, the SiV containing FNDs can be used to target the cell cytoplasm.

Cell viability studies demonstrated that at the tested concentrations of FNDs, CHO-K1 cells show good viability after 4h and even after 24h (Figure S2). A t-test conclusively shows that fluctuations in cell viability are not statistically significant, meaning that the FNDs are non-toxic to the cells at the tested concentrations.

Biological imaging studies were performed using Chinese Hamster Ovarian cell line (CHO-K1 cells) that was incubated with 50 μg/mL of FNDs solutions in cell media (37°C + 5% CO_2_). These were subsequently fixed with a 4% paraformaldehyde/PBS solution, stained with a nucleus stain, NucBlue, and subsequently imaged using an A1 Nikon confocal scanning laser microscope at room temperature equipped with 405 nm, 488 nm and 561 nm excitation laser sources.

The two types of FNDs – namely the ones containing the NV centers and the ones containing the SiV centers, can be individually distinguished based on their spectral differences (Figure S3). Our results show that FND particles readily internalize into the cells with no apparent toxicity (Figure S3) and can be visualized using a standard confocal microscopy set-up. We are able to track FND probes after several hours that helped in understanding complex biological processes happening inside cells upon nanoparticles internalization.

**Figure 3** (a-d) shows the efficient uptake of FNDs hosting SiV color centers into the cells after 3 hours of incubation. Confocal microscopy revealed that the internalization is relatively spontaneous, without additional chemical or mechanical force. Additionally, upon comparing confocal images of control and cells containing FNDs (Figure 3) we observed that the cell nucleus appears to remain healthy and undamaged following the introduction of FNDs. Previous studies by others have demonstrated that internalization of FNDs occurs through endocytosis, specifically via macropinocytosis.^[26]^ This is the most likely uptake for the SiV containing FNDs while FNDs hosting NV color centers would be internalized via a different non-receptor mediated process as their surface is conjugated with the TAT peptide (Figure 3 (e-h)). It can be seen that SiV FNDs appear to spread over the entire cell cytoplasm (Figure 3(a-d)), while NV FNDs with TAT peptide localized to discrete areas (Figure 3 (e-h)).

**Figure 3.**
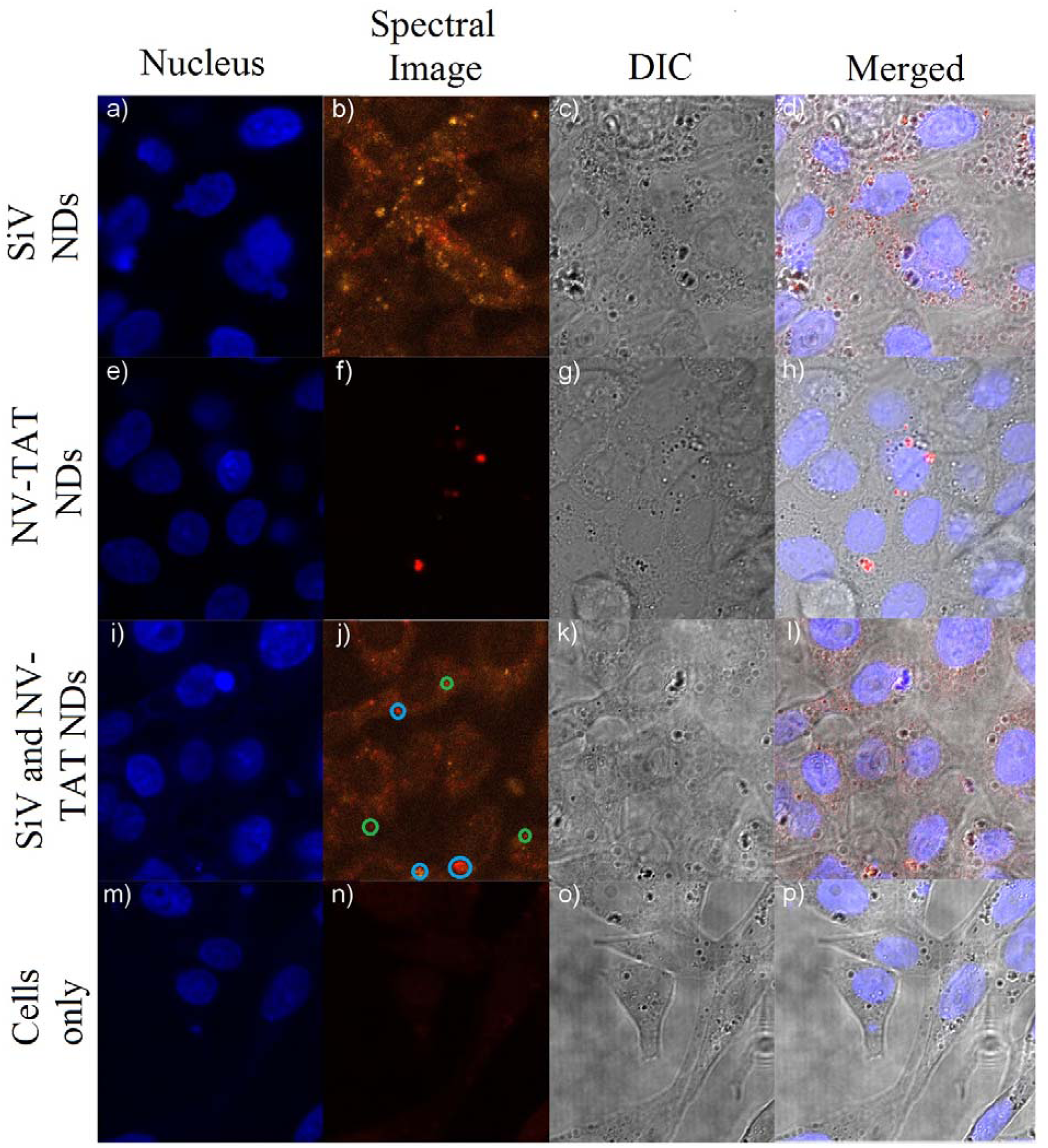
Confocal microscopy of fixed CHO-K1 cells. Cells containing FNDs hosting SiV (a-d), NV (e-h), both SiV and NV-TAT (i-l) and control CHO-K1 cells (m-p) were imaged using confocal microscopy. 405 nm excitation (a, e, i, m) shows the NucBlue stained nucleus of the cells. 561 nm excitation through a spectrometer (b, f, l, n) shows the bright emission from the various FNDs color centers (localized red spheres). The NV (blue circles) and SiV (green circles) color centers are labelled for clarity. The NV emitters preferentially accumulate at the periphery of the nucleus, while the SiV emitters are dispersed throughout the cell cytoplasm. A wide field image (c, g, k, o) and merged (d, h, l, p) image shows the scanned area of interest.

The ND-TAT particles in conjunction with SiV containing FNDs allow the dual tagging of CHO-K1 cells. Figure 3 (i-l) show successful dual tagging of the CHO-K1 cells containing FNDs hosting either SiV or NV color centers. We show the FNDs localization to different areas of the cell sample, the NV emitters (blue circles) preferentially accumulate at the periphery of the nucleus, while the SiV emitters (green circles) are dispersed throughout the cell cytoplasm. Figure 3 (m-p) show the control, non-treated CHO-K1 cells with no fluorescence. To determine unambiguously the positions of the FNDs, a representative 3D constructed confocal z-stack is shown in **Figure 4**, where individual images taken at varying heights are spliced together to create a 3D rendering of the sample. The obtained z-stack images confirm that the FNDs are being internalized by the cells, with NV FNDs-TAT accumulate in close proximity to nucleus, but are not entering the nucleus of the cells. This shows that the FND particles can be used as dual tags in a cellular environment with no significant damage observed.

**Figure 4.**
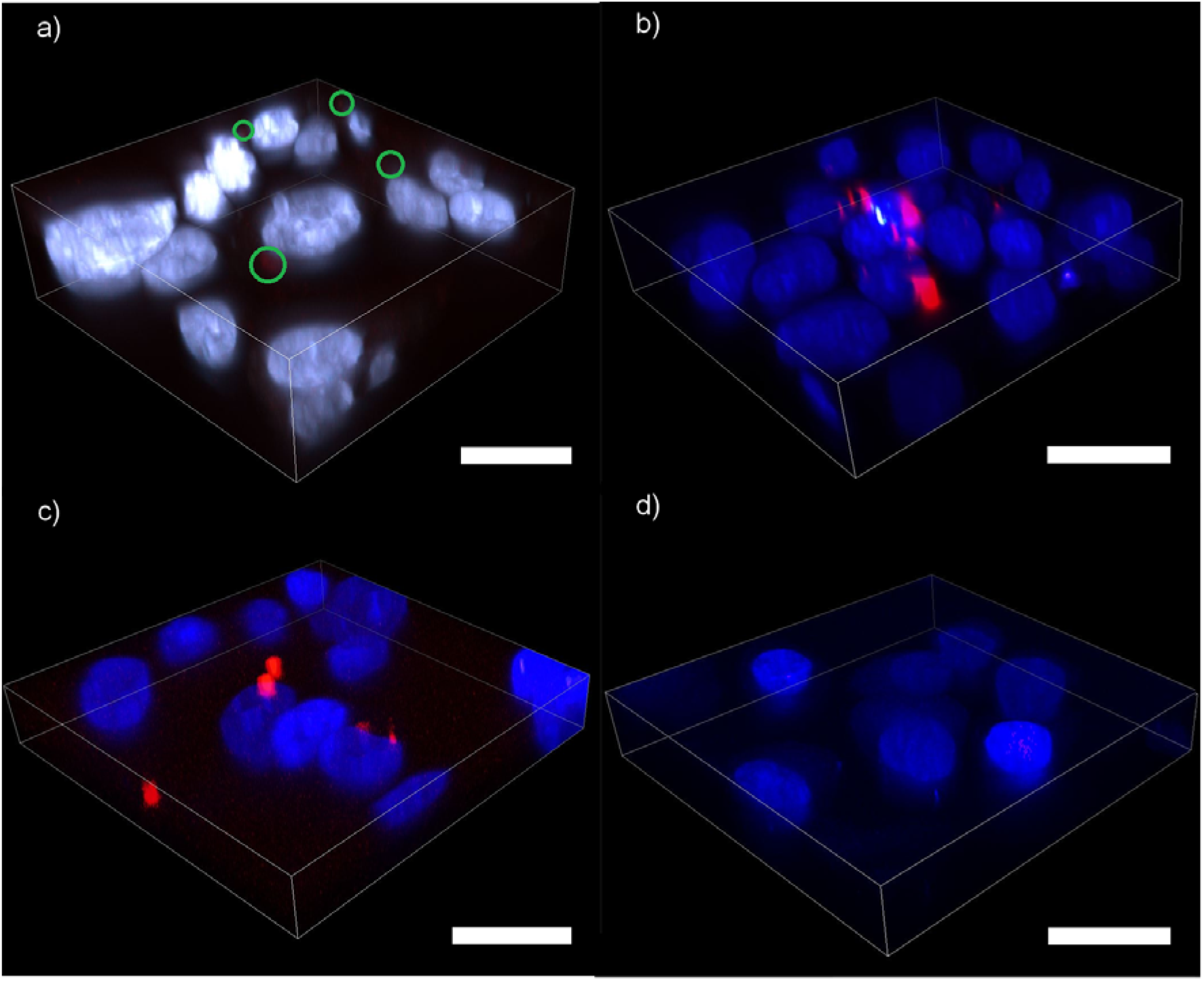
3D z-stack of CHO-K1 cells hosting FNDs. 3D constructed z-stack images of CHOK1 cells hosting (a) SiV (b) NV (c) SiV and NV-TAT containing FNDs and (d) control cells were imaged under 561 nm excitation to reduce cell auto fluorescence and maximise the collected signal from the FNDs particles. Scale bar 20 μm.

The efficiency of uptake for the FNDs particles into the CHO-K1 cells was also studied. A 50 μL solution of FNDs containing either NV or SiV color centers was introduced into the cultured CHO-K1 cells as described previously, with the cells then subsequently fixed and stained after 5, 30, 60, 180 and 300 minutes of incubation at 37°C. **Figure 5** (a-e) shows confocal images of the cells after the various times of incubation. After 5 minutes of incubation, Figure 5a show individual FNDs dispersed throughout the cell interior while a small number of FNDs have been successfully internalized by the CHO-K1 cells. Figure 5 (a-e) shows that the FNDs particles are efficiently uptaken into the cells after 2 hours (between 1 and 3 hours of incubation). The FNDs are likely internalized via endocytic pathways, where the particles are engulfed, then suspended within small vesicles (endosomes). It has been shown for other cell lines that the FNDs are usually trapped in endosomes for an hour prior to translocation to the cytoplasm following degradation of the vesicles.^[26]^ This did not appear to be the case here, as the FNDs appear to accumulate in cellular vesicles. As the time increases from 5 to 300 minutes, we further observe that the ND particles are agglomerating inside the cell. Specifically, we detect that after 300 minutes, the FNDs form large aggregates, likely to be held within lysosomes inside the CHO-K1 cells, located in close proximity to the nucleus. Controlled agglomeration of nanoparticles can lead to novel applications in drug delivery applications inclusive of controlled and sustained release of a drug into the biological system.

**Figure 5.**
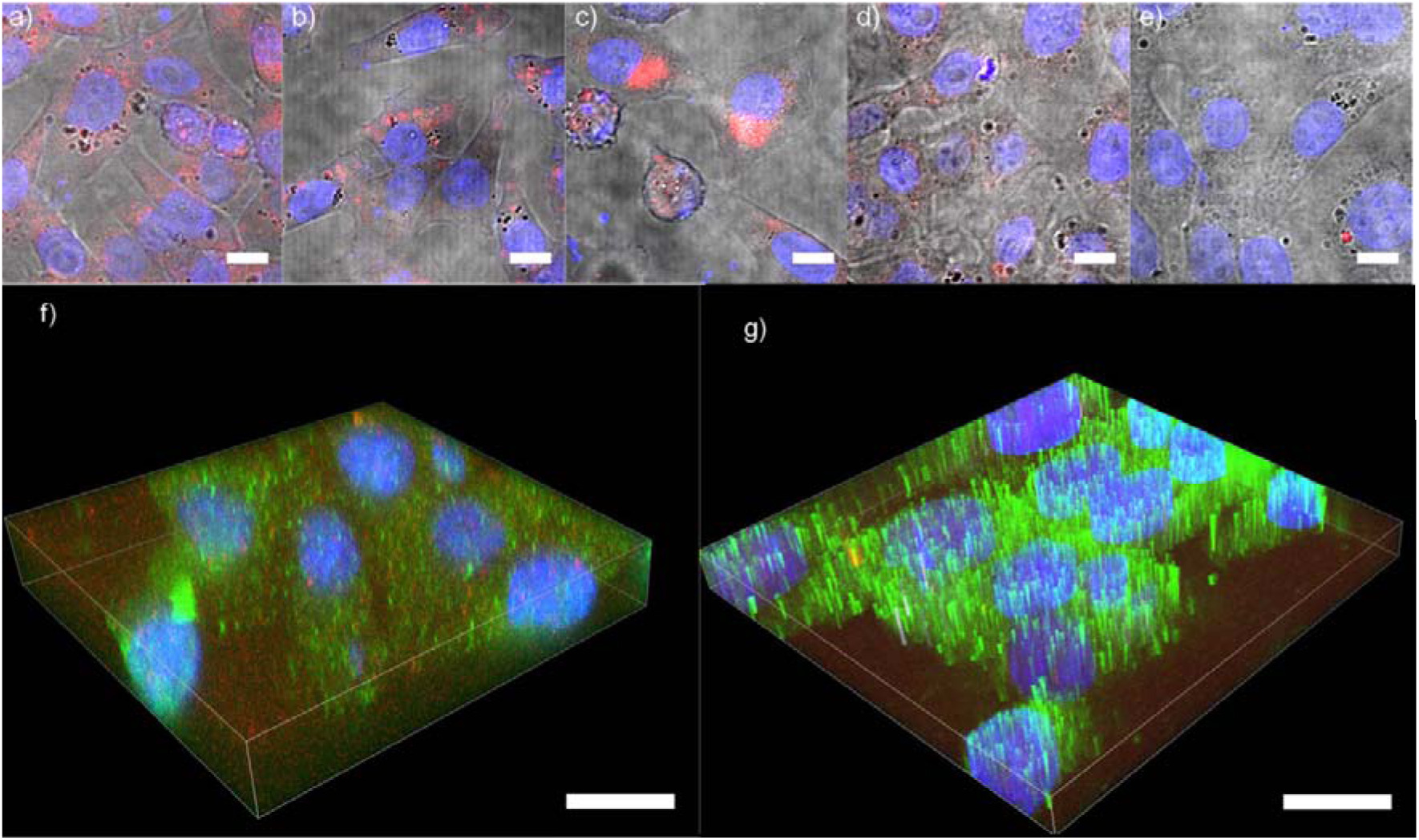
Uptake time study for FNDs hosting NV and SiV color centers. The cells hosting the FNDs particles were fixed after (a) 5 min, (b) 30 min, (c) 60 min, (d) 180 min and (e) 300 min, they were imaged with a 561 nm excitation source. The FNDs are initially dispersed but over time agglomerate near the cell nucleus. Scale bar 10 μm. Co-localization study of FNDs against primary LAMP1 (f) and RAB7 (g) proteins stained the endosomes or lysosomes and the endoplasmic reticulum, respectively. Secondary anti-mouse Alexa Fluor 488 antibodies were used to visualize primary LAMP1 and RAB7 antibobies. Scale bar 20 μm.

To better determine the localization of the FNDs within the cells, co-localization staining studies were undertaken using antibodies against LAMP1 (Lysosomal-associated membrane protein) and RAB7 protein. LAMP1 primarily stains the late endosomes or lysosomes while the RAB7 targets the endoplasmic reticulum (Figure 5(f-g)). The co-localization studies show (**Table 1**) that the FND fluorescence co-incides with LAMP1 fluorescence, supporting the hypothesis that the FNDs are held within lysosomes. It is a known phenomenon that some types of nanoparticles induce a process of autophagy.^[27-29]^ Typically, in autophagy process inside stressed cells is induced to degrade unnecessary or dysfunctional cellular components that can provide energy. This process is suppressed under normal physiological conditions. When external materials, such as nanoparticles, are introduced into cells, cells can induce autophagy pathways to eliminate these extraneous particles. This however requires further studies in order to confirm the fate of these particles. Despite that, localization of FNDs can be further improved by attaching more specific targeting molecules, such as antibodies.

**Table 1.** Co-localization test for control and stained cells. The LAMP1 shows co-localization as the Pearson’s R value tends towards 1.

| Sample | Pearson's R<br>(no<br>Threshold) | Pearson's R<br>(below/above<br>Threshold) | Manders'<br>M1/M2 | Costes P-Value | Li's ICQ |
| --- | --- | --- | --- | --- | --- |
| Cells only (Red vs<br>Green) | 0.00 | -0.00/-0.04 | 0.01/0.27 | 0.98 | 0.112 |
| LAMP1 (Red vs<br>Green) | 0.82 | 0.70/0.15 | 1.00/1.00 | 1.00 | 0.312 |
| Rab7 (Red vs<br>Green) | 0.10 | 0.00/-0.57 | 1.00/1.00 | 1.00 | 0.071 |

Co-localization or degree of overlap of two images, between the ND fluorescence and the cells is determined through a number of statistical parameters. The confidence of co-localization between two color channels, such as the green LAMP1 or RAB7 stained channel and NIR FND channel, increases as the Pearson’s coefficient R tends towards 1. An R value of 0 is equivalent to random noise. The Manders Coefficient M1/M2 is a secondary check on the validity of the test and tends towards 1; it describes the degree of overlapping between the red and green pixels between our particles and stain respectively. Similarly, the Costes P-value and Li’s intensity correction quotient (ICQ) values are separate validity checks and tend towards 95% and 0.5, respectively, when there is co-localization between two images. The Pearson’s R value for the LAMP1 and RAB7 is 0.82 and 0.1 respectively indicating moderate to strong co-localization of intensity distributions between the LAMP1 and FND particles, while there is little to no co-localization between the FNDs and RAB7. The Manders’ M1 and M2 values indicate significant overlap between the red and green pixels for our samples, except for the cells only control sample, as expected. Additional confirmation of co-localization is Li’s ICQ, which indicates moderate co-localization between the LAMP1 stain and FNDs (0.312) and very low for the RAB7 staining and cell only samples (0.07 and 0.32, respectively). By employing statistical analysis, we are able to show that the FNDs localize to the lysosome organelles in the CHO-K1 cells due to moderate to strong co-localization (Table 1).

In conclusion, we demonstrated large scale production of FNDs containing NIR emitters (SiV) suitable for biological labeling. We further show convincing evidence for co-localized uptake of FNDs containing SiV or NV color centers into CHO-K1 cells with no apparent toxicity and can be visualized using a standard confocal set-up. Our uptake timing experiments showed cell-induced aggregation of FNDs over time that most likely correlates to the autophagy process inside cells. Finally, co-localization study demonstrates that after a prolong incubation the FNDs located inside late endosomes/lysosome, but localization of FNDs can be further improved by attaching more specific targeting molecules, such as antibodies. Our results provide an important stepping stone for the effective use of FNDs for heavily sought after bio-sensing, drug delivery and bio-imaging applications, showing that FNDs are a promising biomedical research tool.

## Supporting Information

Supporting Information is available from the Wiley Online Library or from the author.

## Acknowledgements

We would like to acknowledge Michael Johnson and Robert Wooley for their assistance with the confocal microscopy, and Mika Westerhausen for useful discussions. O.S. acknowledges NHMRC-ARC Dementia research development fellowship (APP1101258). I.A. is the recipient of an Australian Research Council Discovery Early Career Research Award (Project No. DE130100592).

---

**Fluorescent nanodiamonds (FNDs) hosting the silicon vacancy (SiV) emitters** that are suitable for bio-imaging are successfully fabricated on a large scale to produce NIR luminescent probes. Simultaneous and targeted labeling of different regions of cell interior using two different types of FNDs, such as containing SiV and NV defects, is also demonstrated to investigate intercellular processes.

**ToC figure**

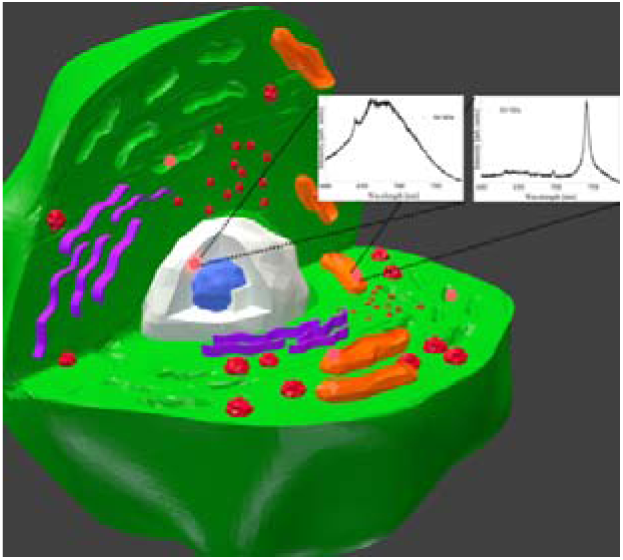

